# Clinical benefit of remdesivir in rhesus macaques infected with SARS-CoV-2

**DOI:** 10.1101/2020.04.15.043166

**Authors:** Brandi N. Williamson, Friederike Feldmann, Benjamin Schwarz, Kimberly Meade-White, Danielle P. Porter, Jonathan Schulz, Neeltje van Doremalen, Ian Leighton, Claude Kwe Yinda, Lizzette Pérez-Pérez, Atsushi Okumura, Jamie Lovaglio, Patrick W. Hanley, Greg Saturday, Catharine M. Bosio, Sarah Anzick, Kent Barbian, Tomas Cihlar, Craig Martens, Dana P. Scott, Vincent J. Munster, Emmie de Wit

## Abstract

**Background:** Effective therapeutics to treat COVID-19 are urgently needed. Remdesivir is a nucleotide prodrug with in vitro and in vivo efficacy against coronaviruses. Here, we tested the efficacy of remdesivir treatment in a rhesus macaque model of SARS-CoV-2 infection.

**Methods:** To evaluate the effect of remdesivir treatment on SARS-CoV-2 disease outcome, we used the recently established rhesus macaque model of SARS-CoV-2 infection that results in transient lower respiratory tract disease. Two groups of six rhesus macaques were infected with SARS-CoV-2 and treated with intravenous remdesivir or an equal volume of vehicle solution once daily. Clinical, virological and histological parameters were assessed regularly during the study and at necropsy to determine treatment efficacy.

**Results:** In contrast to vehicle-treated animals, animals treated with remdesivir did not show signs of respiratory disease and had reduced pulmonary infiltrates on radiographs. Virus titers in bronchoalveolar lavages were significantly reduced as early as 12hrs after the first treatment was administered. At necropsy on day 7 after inoculation, lung viral loads of remdesivir-treated animals were significantly lower and there was a clear reduction in damage to the lung tissue.

**Conclusions:** Therapeutic remdesivir treatment initiated early during infection has a clear clinical benefit in SARS-CoV-2-infected rhesus macaques. These data support early remdesivir treatment initiation in COVID-19 patients to prevent progression to severe pneumonia.

## Introduction

Effective treatments for COVID-19 are urgently needed. While a large number of investigational as well as approved and repurposed drugs have been suggested to have utility for treatment of COVID-19, preclinical data from animal models can guide a more focused search for effective treatments in humans by ruling out treatments without proven efficacy in vivo. Remdesivir (GS-5734) is a nucleotide analog prodrug with broad antiviral activity^1^, including against coronaviruses^2^, that is currently investigated in COVID-19 clinical trials worldwide, including in China, the US and Europe (summarized in^3^). In animal models, remdesivir treatment was effective against MERS-CoV and SARS-CoV infection.^2,4,5^ In vitro, remdesivir inhibited replication of SARS-CoV-2.^6,7^ Moreover, in vitro experiments have shown that mutations conferring resistance to remdesivir do not easily emerge in coronaviruses^8^. Here, we investigated the efficacy of remdesivir treatment in our recently established rhesus macaque model of SARS-CoV-2 infection. In this model, infected rhesus macaques develop mild to moderate, transient respiratory disease with pulmonary infiltrates visible on radiographs, and a shedding pattern similar to that observed in COVID-19 patients^9^. Therapeutic treatment of rhesus macaques with remdesivir shortly before the peak of virus replication resulted in a significant clinical improvement, reduction in pulmonary infiltrates, and a reduction in pulmonary pathology.

## Methods

### Ethics and biosafety statement

All animal experiments were approved by the Institutional Animal Care and Use Committee of Rocky Mountain Laboratories, NIH and carried out by certified staff in an Association for Assessment and Accreditation of Laboratory Animal Care (AAALAC) International accredited facility, according to the institution’s guidelines for animal use, following the guidelines and basic principles in the NIH Guide for the Care and Use of Laboratory Animals, the Animal Welfare Act, United States Department of Agriculture and the United States Public Health Service Policy on Humane Care and Use of Laboratory Animals. Rhesus macaques were housed in adjacent individual primate cages allowing social interactions, in a climate-controlled room with a fixed light-dark cycle (12-hr light/12-hr dark). Animals were monitored at least twice daily throughout the experiment. Commercial monkey chow, treats, and fruit were provided twice daily by trained personnel. Water was available ad libitum. Environmental enrichment consisted of a variety of human interaction, manipulanda, commercial toys, videos, and music. The Institutional Biosafety Committee (IBC) approved work with infectious SARS-CoV-2 strains under BSL3 conditions. Sample inactivation was performed according to IBC-approved standard operating procedures for removal of specimens from high containment.

### Study design

To evaluate the effect of remdesivir treatment on SARS-CoV-2 disease outcome, we used the recently established rhesus macaque model of SARS-CoV-2 infection that results in transient lower respiratory tract disease^9^. Twelve animals were randomly assigned to two groups and inoculated as described previously with a total dose of 2.6×10^6^ TCID50 of SARS-CoV-2 strain nCoV-WA1-2020 via intranasal, oral, ocular and intratracheal routes. The efficacy of therapeutic remdesivir treatment was tested in two groups of six adult rhesus macaques (3 males and 3 females each; 3.6-5.7kg). Due to the acute nature of the SARS-CoV-2 model in rhesus macaques, therapeutic treatment was initiated at 12 hours after inoculation with SARS-CoV-2 and continued once daily through 6 days post inoculation (dpi). One group of rhesus macaques was treated with a loading dose of 10mg/kg remdesivir, followed by a daily maintenance dose of 5 mg/kg. The other group of six animals served as infected controls and were administered an equal dose volume (i.e. 2 ml/kg loading dose and 1 ml/kg thereafter) of vehicle solution (12% sulfobutylether-β-cyclodextrin in water and hydrochloric acid, pH3.5) according to the same treatment schedule. This dosing scheme in rhesus macaques mimics the daily dosing tested in clinicals studies with COVID-19 patients and results in a similar systemic drug exposure. Treatment was delivered as an intravenous bolus injection (total dose delivered over approximately 5 minutes) administered alternatingly in the left or right cephalic or saphenous veins. The animals were observed twice daily for clinical signs of disease using a standardized scoring sheet as described previously^10^; the same person, who was blinded to the group assignment of the animals, assessed the animals throughout the study. The predetermined endpoint for this experiment was 7 dpi. Nose, throat and rectal swabs were collected daily during treatment administration. Clinical exams were performed on 0, 1, 3, 5, and 7 dpi on anaesthetized animals. On exam days, clinical parameters such as bodyweight, temperature, pulse oximetry, blood pressure and respiration rate were collected, as well as dorsal-ventral and lateral chest radiographs. Radiographs were analyzed by a clinical veterinarian blinded to the group assignment of the animals. On 1, 3 and 7 dpi a bronchoalveolar lavage (BAL) was performed using 10ml of sterile saline. After euthanasia on 7 dpi, necropsies were performed. The percentage of gross lung lesions were scored by a board-certified veterinary pathologist blinded to the group assignment of the animals and samples of the following tissues were collected: cervical lymph node, conjunctiva, nasal mucosa, oropharynx, tonsil, trachea, all lung lobes, mediastinal lymph node, right and left bronchus, heart, liver, spleen, kidney, stomach, duodenum, jejunum, ileum, cecum, colon, and urinary bladder. Histopathological analysis of tissue slides was performed by a board-certified veterinary pathologist blinded to the group assignment of the animals.

### Virus and cells

SARS-CoV-2 isolate nCoV-WA1-2020 (MN985325.1)^11^ (Vero passage 3) was kindly provided by CDC and propagated once in Vero E6 cells in DMEM (Sigma) supplemented with 2% fetal bovine serum (Gibco), 1 mM L-glutamine (Gibco), 50 U/ml penicillin and 50 μg/ml streptomycin (Gibco) (virus isolation medium). The virus stock used was 100% identical to the initial deposited Genbank sequence (MN985325.1) and no contaminants were detected. VeroE6 cells were maintained in DMEM supplemented with 10% fetal calf serum, 1 mM L-glutamine, 50 U/ml penicillin and 50 μg/ml streptomycin.

### Remdesivir (GS-5734)

Remdesivir (RDV; GS-5734) was manufactured at Gilead Sciences by the Department of Process Chemistry (Alberta, Canada) under Good Manufacturing Practice (GMP) conditions. Batch number 5734-BC-1P was solubilized in 12% sulfobutylether-β-cyclodextrin in water and matching vehicle solution was provided to NIH.

### Liquid chromatography mass spectrometry

Tributylamine was purchased from Millipore Sigma. LCMS grade water, acetone, methanol, isopropanol and acetic acid were purchased through Fisher Scientific. All synthetic standards for molecular analysis were provided by Gilead Sciences Inc. Serum and cleared lung homogenates were gamma-irradiated (2 MRad) to inactivate infectious virus potentially present in these samples prior to analysis. Samples were prepared for small molecule analysis by diluting a 50 µL aliquot of either serum or clarified lung homogenate with 950 µL of 50% acetone, 35% methanol, 15% water (v/v) on ice. Samples were incubated at room temperature for 15 min and then centrifuged at 16k xg for 5 minutes. The clarified supernatants (850 µL) were recovered and taken to dryness in a Savant™ DNA120 SpeedVac™ concentrator (Thermo Fisher). Samples were resuspended in 100 µL of 50% methanol, 50% water (v/v) and centrifuged as before. The supernatant was taken to a sample vial for LCMS analysis. Samples were separated using an ion-pairing liquid chromatography strategy on a Sciex ExionLC™ AC system. Samples were injected onto a Waters Atlantis T3 column (100 Å, 3 µm, 3 mm X 100 mm) and eluted using a binary gradient from 5 mM tributylamine, 5 mM acetic acid in 2% isopropanol, 5% methanol, 93% water (v/v) to 100% isopropanol over 5.5 minutes. Analytes were measured using a Sciex 5500 QTRAP® mass spectrometer in negative mode. Multiple reaction monitoring was performed using two signal pairs for each analyte and signal fidelity was confirmed by collecting triggered product ion spectra and comparing back to spectra of synthetically pure standards.

All analytes were quantified against an 8-point calibration curve of the respective synthetic standard prepared in the target matrix (i.e. serum or cleared lung homogenate) and processed in the same manner as experimental samples. Limit of quantification (LOQ) was approximated at a signal to noise of 10. The LOQs for the measured molecules in each matrix were 5 nM for GS-441524 in both lung homogenate and serum, 1 nM for GS-704277 in both lung homogenate and serum and 0.08 nM for GS-5734 in serum. Instability of GS-5734 and the tri-phosphorylated nucleotide metabolite in the lung homogenate during tissue lysis prevented detection of these metabolites in the lung tissue.

### Quantitative PCR

RNA was extracted from swabs and BAL using the QiaAmp Viral RNA kit (Qiagen) according to the manufacturer’s instructions. Tissues (30 mg) were homogenized in RLT buffer and RNA was extracted using the RNeasy kit (Qiagen) according to the manufacturer’s instructions. For detection of viral RNA, 5 µl RNA was used in a one-step real-time RT-PCR E assay^12^ using the Rotor-Gene probe kit (Qiagen) according to instructions of the manufacturer. In each run, standard dilutions of RNA standards counted by droplet digital PCR were run in parallel, to calculate copy numbers in the samples.

### Virus titration

Virus titrations were performed by end-point titration in Vero E6 cells. Tissue was homogenized in 1ml DMEM using a TissueLyser (Qiagen). Cells were inoculated with 10-fold serial dilutions of swab and BAL samples. Virus isolation was performed on lung tissues by homogenizing the tissue in 1ml DMEM and inoculating Vero E6 cells in a 24 well plate with 250 µl of cleared homogenate and a 1:10 dilution thereof. One hour after inoculation of cells, the inoculum was removed and replaced with 100 µl (virus titration) or 500 µl virus isolation medium. Six days after inoculation, CPE was scored and the TCID50 was calculated.

### Histopathology and immunohistochemistry

Histopathology and immunohistochemistry were performed on rhesus macaque tissues. After fixation for a minimum of 7 days in 10% neutral-buffered formalin and embedding in paraffin, tissue sections were stained with hematoxylin and eosin (HE). To detect SARS-CoV-2 antigen, immunohistochemistry was performed using a custom-made rabbit antiserum against SARS-CoV-2 N at a 1:1000 dilution. Stained slides were analyzed by a board-certified veterinary pathologist.

### Next generation sequencing of viral RNA

Viral RNA was extracted as described above. cDNAs were prepared according to Briese et al., with minor modifications^13^. Briefly, 3 to 12 µl of extracted RNA was depleted of rRNA using Ribo-Zero Gold H/M/R (Illumina) and then reverse-transcribed using random hexamers and SuperScript IV (ThermoFisher Scientific). Following RNaseH treatment, second strand synthesis was performed using Klenow fragment (New England Biolabs) and resulting double-stranded cDNAs were treated with a combined mixture of RiboShredder RNase Blend (Lucigen) and RNase, DNase-free, high conc (Roche Diagnostics, Indianapolis, IN) and then purified using Ampure XP bead purification (Beckman Coulter). Kapa’s HyperPlus library preparation kit (Roche Sequencing Solutions) was used to prepare sequencing libraries from the double-stranded cDNAs. To facilitate multiplexing, adapter ligation was performed with KAPA Unique Dual-Indexed Adapters and samples were enriched for adapter-ligated product using KAPA HiFi HotStart Ready mix and 7 PCR amplification cycles, according to the manufacturer’s manual. Pools consisting of eight sample libraries were used for hybrid-capture virus enrichment using myBaits® Expert Virus SARS-CoV-2 panel and following the manufacturer’s manual, version 4.01, with 14 cycles of post-capture PCR amplification (Arbor Biosciences). Purified, enriched libraries were quantified using Kapa Library Quantification kit (Roche Sequencing Solutions) and sequenced as 2 × 150 base pair reads on the Illumina NextSeq 550 instrument (Illumina).

Raw fastq reads were trimmed of Illumina adapter sequences using cutadapt version 1.12^14^ and then trimmed and filtered for quality using the FASTX-Toolkit (Hannon Lab). Remaining reads were mapped to the SARS-CoV-2 2019-nCoV/USA-WA1/2020 genome (MN985325.1) using Bowtie2 version 2.2.9^15^ with parameters --local --no-mixed -X 1500. PCR duplicates were removed using picard MarkDuplicates (Broad Institute) and variants were called using GATK HaplotypeCaller version 4.1.2.0^16^ with parameter - ploidy 2. Variants were filtered for QUAL > 1000 and DP > 20 using bcftools.

### Statistical analysis

Statistical analyses were performed using GraphPad Prism software version 8.2.1.

### Data sharing

All data included in this manuscript have been deposited in Figshare (https://doi.org/10.35092/yhjc.12111570).

## Results

### Remdesivir is distributed to the main target tissue of SARS-CoV-2, the lungs

Two groups of six rhesus macaques were inoculated with SARS-CoV-2 strain nCoV-WA1-2020. Twelve hours post inoculation, one group was administered 10mg/kg intravenous remdesivir and the other group was treated with an equal volume of vehicle solution (2ml/kg). Treatment was continued 12hrs after the first treatment, and every 24 hrs thereafter with a dose of 5 mg/kg remdesivir or an equal volume of vehicle solution (1ml/kg). The serum concentration of remdesivir was determined in serum collected 12 hrs after the initial treatment and 24 hrs after subsequent treatments were administered and immediately before the next dose of treatment was administered. Detectable levels of remdesivir (prodrug GS-5734) as well as the downstream alanine metabolite (GS-704277) and parent nucleoside (GS-441524) were observed in all remdesivir-treated animals (Fig. S1A). Serum levels of the prodrug and the downstream metabolites were consistent with previously published plasma levels of these compounds in healthy rhesus macaques, which showed a short systemic half-life for GS-5734 (0.39 hrs) resulting in transient conversion to the intermediate GS-704277 and persistence of the downstream GS-441524 product at higher plasma levels^17^.

Concentrations of metabolite GS-441524 were determined in lung tissue collected from each lung lobe on 7 dpi, 24 hrs after the last remdesivir treatment was administered and was readily detectable in all remdesivir-treated animals. GS-441524 was generally distributed amongst all six lobes of the lung (Fig S1B). GS-704277 was not detected in the lung tissue. While the pharmacologically active metabolite of remdesivir is the triphosphate of GS-441524, lung homogenate samples spiked with the triphosphate metabolite demonstrated rapid decay of the metabolite in this matrix (data not shown). GS-441524 levels were taken as a surrogate for tissue loading and suggest that the current dosing strategy delivered drug metabolites to the sites of SARS-CoV-2 replication in infected animals.

### Lack of respiratory disease in rhesus macaques infected with SARS-CoV-2 and treated with remdesivir

After inoculation with SARS-CoV-2, the animals were assigned a daily clinical score based on a pre-established scoring sheet in a blinded fashion. Twelve hours after the first remdesivir treatment, clinical scores in remdesivir-treated animals were significantly lower than in control animals receiving vehicle solution. This difference in clinical score was maintained throughout the study (Fig. 1A). Only one of the six remdesivir-treated animals showed mild dyspnea, whereas tachypnea and dyspnea were observed in all vehicle-treated controls (Table S1). Radiographic pulmonary infiltrates are one of the hallmarks of COVID-19 in humans. Radiographs taken on 0, 1, 3, 5, and 7 dpi showed significantly less lung lobe involvement and less severe of pulmonary infiltration in animals treated with remdesivir as compared to those receiving vehicle (Fig. 1B and C).

**Figure 1.**
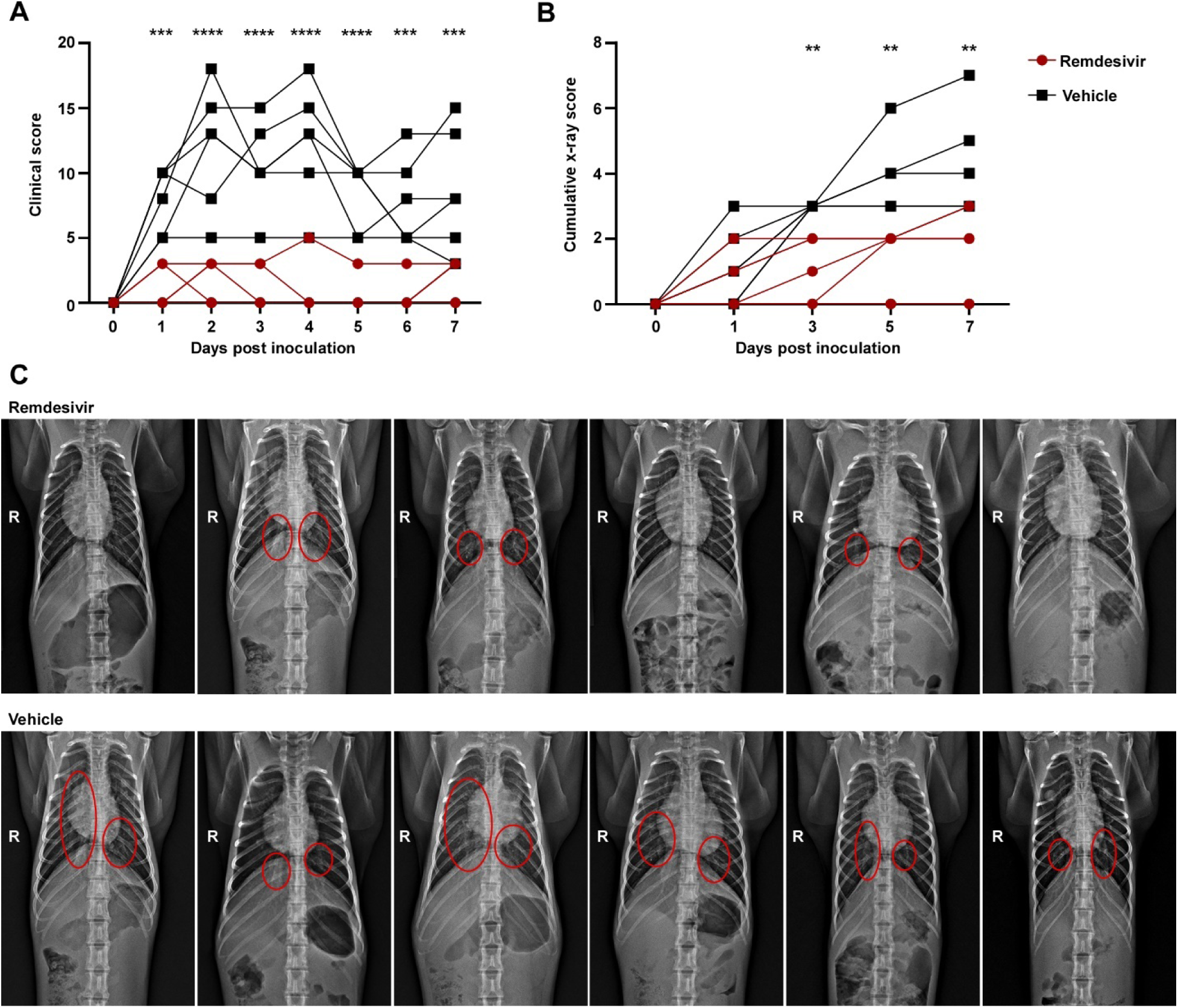
Reduced respiratory disease in rhesus macaques infected with SARS-CoV-2 and treated with remdesivir. Panel A shows daily clinical scores in animals infected with SARS-CoV-2 and treated with remdesivir (red circles) or vehicle solution (black squares). Panel B shows cumulative radiograph scores. Ventro-dorsal and lateral radiographs were scored for the presence of pulmonary infiltrates by a clinical veterinarian according to a standard scoring system (0: normal; 1: mild interstitial pulmonary infiltrates; 2: moderate pulmonary infiltrates perhaps with partial cardiac border effacement and small areas of pulmonary consolidation; 3: severe interstitial infiltrates, large areas of pulmonary consolidation, alveolar patterns and air bronchograms). Individual lobes were scored and scores per animal per day were totaled and displayed. Panel C shows ventro-dorsal radiographs collected from each animal taken on 7 dpi. Areas of pulmonary infiltration are marked with a circle. Statistical analysis was performed using a 2-way ANOVA with Sidak’s multiple comparisons test. ** P<0.01; *** P< 0.001; **** P< 0.0001

### Reduced virus replication in the lower, but not upper respiratory tract after remdesivir treatment

On 1, 3 and 7 dpi BAL were performed as an indicator of virus replication in the lower respiratory tract. Although viral loads in BAL were reduced in remdesivir-treated animals this difference was not statistically significant (Fig. 2A). However, 12 hours after the first remdesivir treatment was administered, the infectious virus titer in BAL was ~100-fold lower in remdesivir-treated animals than controls. By 3 dpi, infectious virus could no longer be detected in BAL from remdesivir-treated animals, whereas virus was still detected in BAL from four out of six control animals (Fig. 2B). Despite this reduction in virus replication in the lower respiratory tract, neither viral loads nor infectious virus titers were reduced in nose, throat or rectal swabs collected from remdesivir-treated animals, except a significant difference in virus titer in throat swabs collected on 1 dpi and in viral loads in throat swabs collected on 4 dpi (Fig. 3).

**Figure 2.**
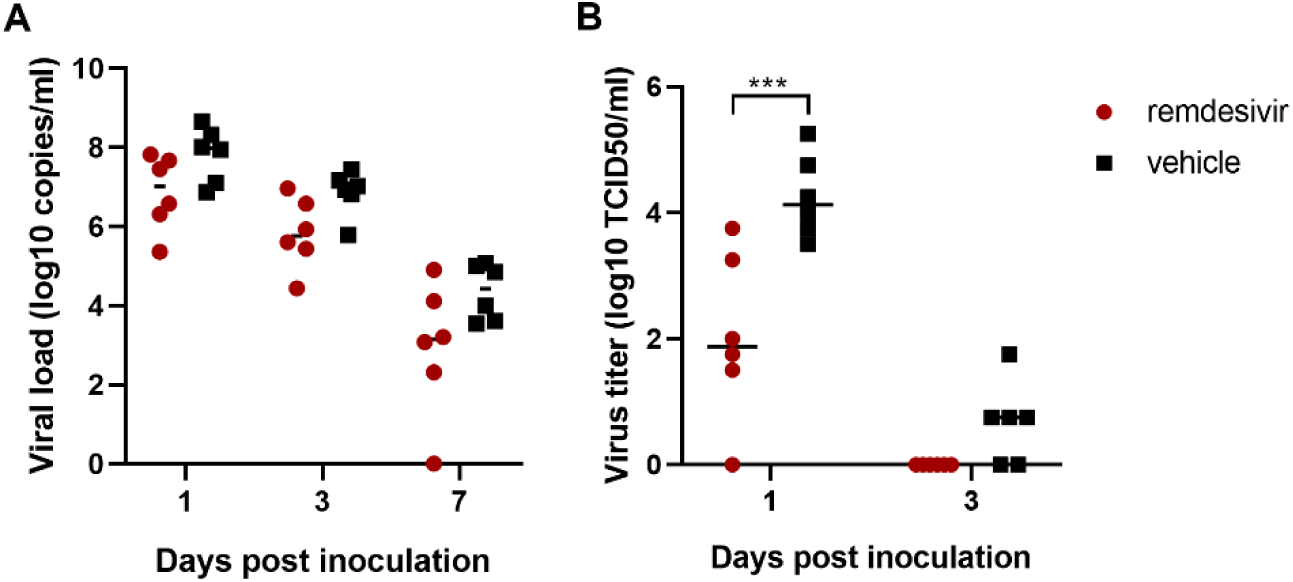
Viral loads and virus titers in bronchoalveolar lavage fluid. Panel A shows viral loads and Panel B shows infectious virus titers in BAL collected from rhesus macaques infected with SARS-CoV-2 and treated with remdesivir (red circles) or vehicle solution (black squares). Statistical analysis was performed using a 2-way ANOVA with Sidak’s multiple comparisons test. *** P< 0.001

**Figure 3.**
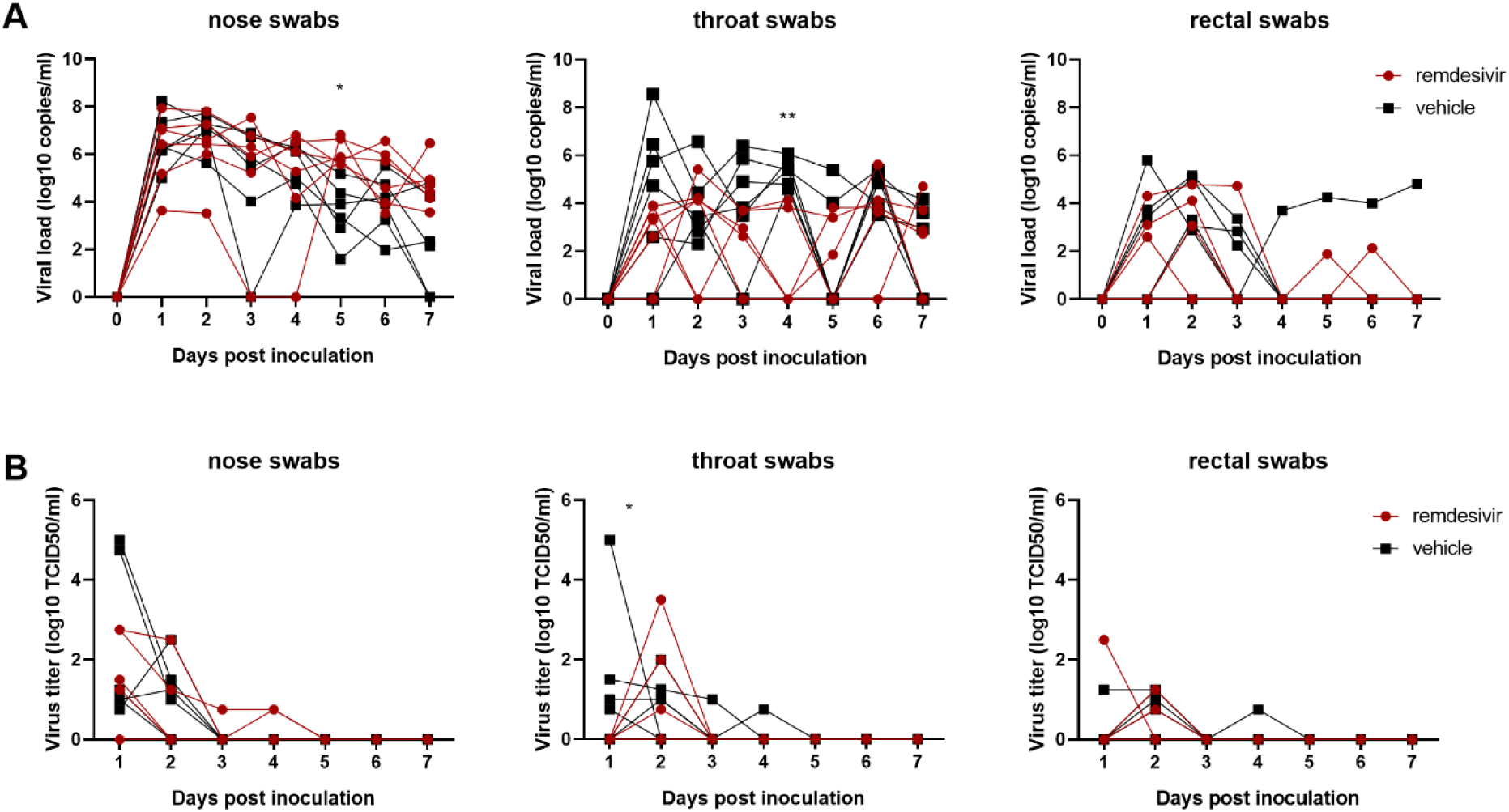
Viral loads and virus titers in swabs collected from rhesus macaques infected with SARS-CoV-2 and treated with remdesivir. Panel A shows viral loads and Panel B shows infectious virus titers in nose, throat and rectal swabs collected daily. Statistical analysis was performed using a 2-way ANOVA with Sidak’s multiple comparisons test. * P<0.05; ** P<0.01

### Decreased viral loads in lungs after remdesivir treatment

All animals were euthanized on 7 dpi. Tissue samples were collected from each lung lobe to compare virus replication in remdesivir-treated and vehicle-treated control animals. In 10 out of 36 lung lobe samples collected from remdesivir-treated animals, viral RNA could not be detected, whereas this was the case in only 3 out of 36 lung lobes collected from control animals. In general, comparison across individual lung lobes in the two groups showed lower geometric mean of viral RNA in remdesivir-treated group (Fig. 4A). Taken together, the viral load was significantly lower in lungs from remdesivir-treated animals than in vehicle-treated controls (Fig. 4B). Virus could be isolated from lung lobes of five out of six vehicle-treated control animals, but none of the lung tissue collected from remdesivir-treated animals was positive in virus isolation. Although fewer tissues from other positions in the respiratory tract were positive by qRT-PCR in remdesivir-treated animals, these differences were not statistically significant (Fig. 4C).

**Figure 4.**
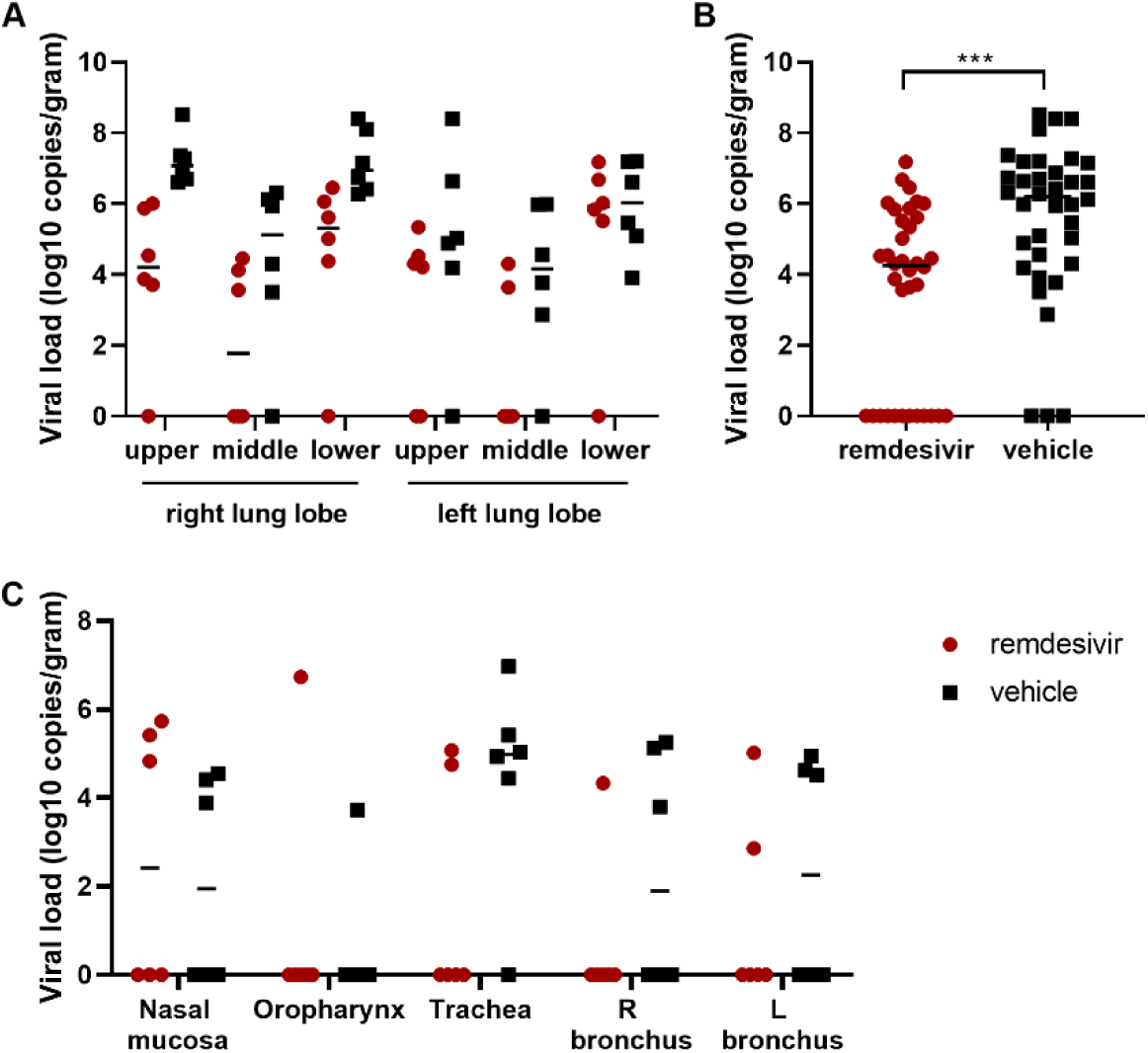
Viral loads in tissues collected from the respiratory tract on 7 dpi. Panel A shows viral loads in all six lung lobes collected from rhesus macaques infected with SARS-CoV-2 and treated with remdesivir (red circles) or vehicle solution (black squares), stratified per lung lobe. In panel B, all viral loads were combined. Statistical analysis was performed using an unpaired t test. ***P<0.001. Panel C shows viral loads in other tissues collected throughout the respiratory tract on 7 dpi.

### Reduced pneumonia after remdesivir treatment

At necropsy on 7 dpi, lungs were assessed grossly for presence of lesions. Gross lung lesions were observed in one out of six remdesivir-treated animals. In contrast, all six vehicle controls had visible lesions, resulting in statistically significantly difference in the area of the lungs affected by lesions (Fig. 5A, B and Fig. 6A, B). This difference was also evident when calculating the lung weight to bodyweight ratio as an indicator of pneumonia, with a statistically significantly lower ratio observed in remdesivir-treated compared to vehicle-treated animals (Fig. 5C). Histologically, there was a clear effect of remdesivir treatment on lung lesions, with fewer and less severe lesions in the remdesivir animals than in vehicle-treated controls. Histologic lung lesions were absent in three of six remdesivir-treated animals; the three remaining animals developed minimal pulmonary pathology. Lesions in these animals were characterized as widely separated, minimal, interstitial pneumonia frequently located in subpleural spaces (Fig. 6C, E). Five out six vehicle-treated animals developed multifocal, mild to moderate, interstitial pneumonia (Fig. 6D, F). Viral antigen was detected in all animals regardless of treatment (Fig. 6G, H).

**Figure 5.**
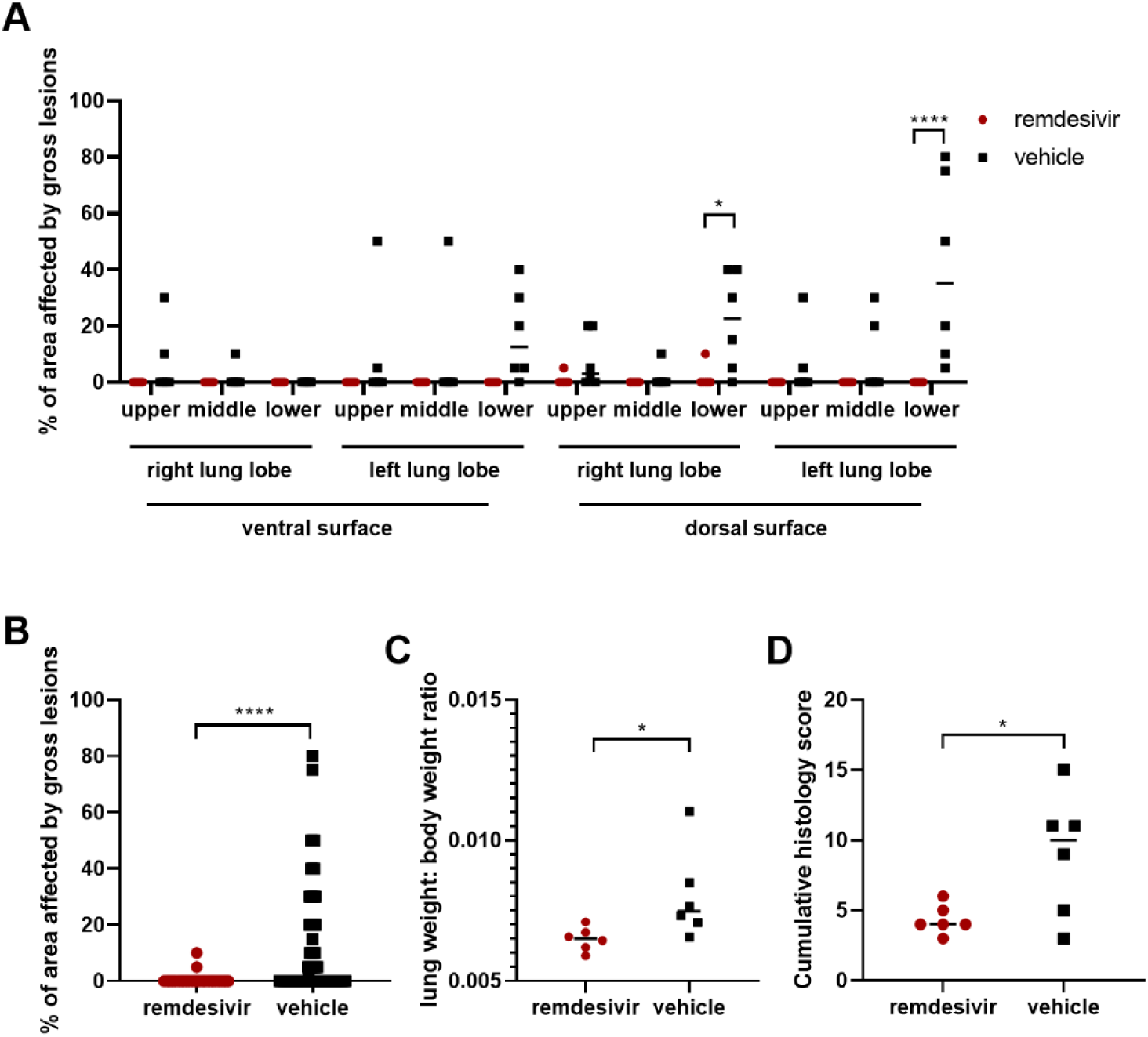
Pathological changes in lungs of rhesus macaques infected with SARS-CoV-2 and treated with remdesivir. Panel A shows the area of each individual lung lobe affected by gross lesions as scored by a veterinary pathologist. In panel B, all data from A are combined. Panel C shows the lung weight: bodyweight ratio as an indicator of pulmonary edema. Panel D shows the cumulative histology score. Each lung lobe was scored for the presence of histologic lung lesions on a predetermined scale (0-4); these values were combined per animal and graphed. Data in panel A were analyzed using a 2-way ANOVA with Sidak’s multiple comparisons test; data in panels B-D were analyzed using an unpaired t test. * P<0.05; **** P<0.0001

**Figure 6.**
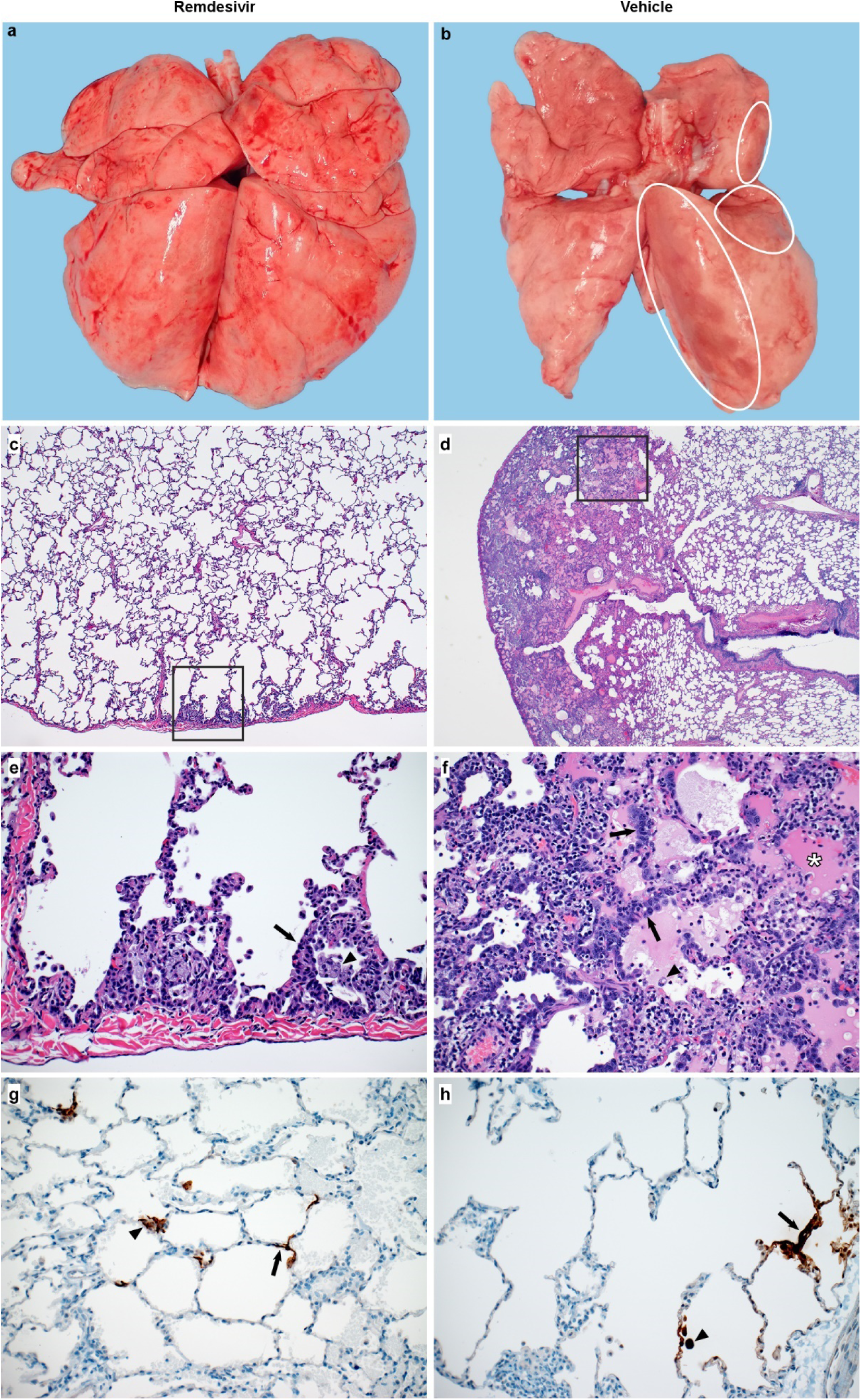
Changes to the lungs of rhesus macaques infected with SARS-CoV-2 and treated with remdesivir. Panel A shows a representative dorsal view of lungs of a remdesivir-treated animal. Panel B shows a representative dorsal view of lungs of a vehicle-treated animal with focally extensive areas of consolidation (circles). Panel C shows minimal subpleural interstitial pneumonia (box) observed in 3 of 6 remdesivir-treated animals. Panel D shows moderate subpleural interstitial pneumonia with edema (box) observed in 5 of 6 vehicle-treated animals. Panel E shows the boxed area from panel C with alveoli lined by type II pneumocytes (arrow) and alveolar spaces containing foamy macrophages (arrowhead). Panel F shows the boxed area from panel E with pulmonary interstitium expanded by edema and moderate numbers of inflammatory cells. Alveoli are lined by type II pneumocytes (arrows). Alveolar spaces are filled with edema (asterisk) and small numbers of pulmonary macrophages (arrowhead). Panel G shows viral antigen in type I pneumocytes (arrow) and type II pneumocytes (arrowhead) of a remdesivir-treated animal. Panel H shows viral antigen in type I pneumocytes (arrow) and macrophage (arrowhead) of a vehicle-treated animal. Magnification C and D: 40x; panel E-H: 200x.

### Absence of resistance mutations

Deep sequencing was successful on samples from all remdesivir-treated animals and vehicle controls. Known mutations in the RNA dependent RNA polymerase that confer resistance to remdesivir in coronaviruses^8^ were not detected in any of the samples tested (Table S2).

## Discussion

Remdesivir is the first antiviral treatment with proven efficacy against SARS-CoV-2 in an animal model of COVID-19. Remdesivir treatment in rhesus macaques infected with SARS-CoV-2 was highly effective in reducing clinical disease and damage to the lungs. The remdesivir dosing used in rhesus macaques is equivalent to that used in humans; however, due to the acute nature of the disease in rhesus macaques, it is hard to directly translate the timing of treatment used to corresponding disease stages in humans. In our study, treatment was administered close to the peak of virus replication in the lungs as indicated by viral loads in bronchoalveolar lavages and the first effects of treatment on clinical signs and virus replication were observed within 12 hours. The efficacy of direct-acting antivirals against acute viral respiratory tract infections typically decreases with delays in treatment initation^18^. Thus, remdesivir treatment in COVID-19 patients should be initiated as early as possible to achieve the maximum treatment effect.

Despite the lack of obvious respiratory signs and reduced virus replication in the lungs of remdesivir-treated animals, there was no reduction in virus shedding. This finding is of great significance for patient management, where a clinical improvement should not be interpreted as a lack of infectiousness. While our study demonstrates the presence of remdesivir metabolites in the lower respiratory tract, the drug levels in upper respiratory tract have not been characterized and novel formulations with alternative route of drug delivery should be considered to improve the distribution to the upper respiratory tract, thereby reducing shedding and the potential transmission risk. However, since severe COVID-19 disease is a result of virus infection of the lungs, this organ is the main target of remdesivir treatment. The bioavailability and protective effect of remdesivir in the lungs of infected rhesus macaques supports treatment of COVID-19 patients with remdesivir. Data from clinical trials in humans are pending, but our data in rhesus macaques indicate that remdesivir treatment should be considered as early as clinically possible to prevent progression to severe pneumonia in COVID-19 patients.

## Acknowledgements

The authors would like to thank Elaine Bunyan (Gilead Sciences) for preparing remdesivir; Darius Babusis (Gilead Sciences) for providing synthetic standards for molecular analysis; Anita Mora (NIAID) for preparing figures; Tina Thomas, Rebecca Rosenke and Dan Long (all NIAID) for assistance with histology; Myndi Holbrook (NIAID) for technical assistance and RMVB staff (NIAID) for animal care. This study was supported by the Intramural Research Program of NIAID, NIH.

