## supplemental materials for "Clinical benefit of remdesivir in rhesus macaques infected with SARS-CoV-2"

**Table S1. Clinical and pathological observations in rhesus macaques inoculated with SARS-CoV-2 and treated with remdesivir.**

| Treatment | Animal | Clinical observations | Observations at necropsy |
|---|---|---|---|
| **Remdesivir** | RM1 | Slightly decreased appetite | Mediastinal lymph nodes enlarged |
| | RM2 | Slightly decreased appetite | None |
| | RM3 | Slightly decreased appetite, pale appearance | Mediastinal lymph nodes enlarged |
| | RM4 | Slightly decreased appetite, slightly dehydrated | Mediastinal lymph nodes enlarged |
| | RM5 | Slightly decreased appetite | Mediastinal lymph nodes enlarged |
| | RM6 | Mild dyspnea, pale appearance | Gross lung lesions; mediastinal lymph nodes enlarged |
| **Vehicle solution** | RM7 | Piloerection, hunched posture, tachypnea, dyspnea, decreased appetite | Gross lung lesions; mediastinal lymph nodes enlarged; focal hemorrhage in colon |
| | RM8 | Piloerection, hunched posture, tachypnea, dyspnea, decreased appetite | Gross lung lesions; mediastinal lymph nodes enlarged |
| | RM9 | Piloerection, hunched posture, tachypnea, dyspnea, decreased appetite | Gross lung lesions; mediastinal lymph nodes enlarged |
| | RM10 | Tachypnea, dyspnea, pale appearance, slightly dehydrated | Gross lung lesions; mediastinal lymph nodes enlarged |
| | RM11 | Piloerection, tachypnea, dyspnea, decreased appetite, pale appearance | Gross lung lesions; mediastinal lymph nodes enlarged |
| | RM12 | Piloerection, tachypnea, dyspnea, decreased appetite | Gross lung lesions; mediastinal lymph nodes enlarged; ~5ml fluid in peritoneum |

**Table S2. Deep sequencing results to confirm absence of known resistance mutations to remdesivir.**

The timepoints for sequencing of BAL and swab samples were selected based on positivity in qRT-PCR to reflect the latest possible timepoints where the majority of animals were positive for that given sample type in qRT-PCR.

| Treatment | Animal no. | Sample | Mean sequencing coverage | F476 (nt 14,878-14,880) | V553 (nt 15,109-15,111) |
|---|---|---|---|---|---|
| Remdesivir | RM1 | BAL[a] | 197.27 | no variants | no variants |
| | | LLLL[b] | 21.05 | no variants | no variants |
| | | Nose swab[c] | 138.89 | no variants | no variants |
| | | RLLL[d] | 11.22 | no variants | no variants |
| | RM2 | BAL | 21.7 | no variants | no variants |
| | | LLLL | 32.65 | no variants | no variants |
| | | Nose swab | 187.29 | no variants | no variants |
| | | RLLL | 19.37 | no variants | no variants |
| | RM3 | BAL | 4.76 | ND[f] | ND |
| | | LLLL | 5.43 | ND | ND |
| | | Nose swab | 49.11 | no variants | no variants |
| | | RLLL | 2.08 | no variants | ND |
| | | Rectal swab[e] | 35.47 | no variants | no variants |
| | RM4 | BAL | 421.43 | no variants | no variants |
| | | LLLL | 81.65 | no variants | no variants |
| | | Nose swab | 31.97 | no variants | no variants |
| | | RLLL | 0.05 | ND | ND |
| | | Rectal swab | 0.56 | ND | ND |
| | RM5 | BAL | 25.37 | no variants | no variants |
| | | LLLL | 0.02 | ND | ND |
| | | Nose swab | 25.13 | no variants | no variants |
| | | RLLL | 0.31 | ND | ND |
| | RM6 | BAL | 17.1 | no variants | no variants |
| | | LLLL | 5.39 | no variants | ND |
| | | Nose swab | 25.27 | ND | no variants |
| | | RLLL | 1.25 | ND | no variants |
| | | Rectal swab | 0.42 | ND | ND |
| Vehicle | RM7 | BAL | 352.51 | no variants | no variants |
| | | LLLL | 123.92 | no variants | no variants |
| | | Nose swab | 1.11 | ND | no variants |
| | | RLLL | 229.79 | no variants | no variants |
| | | Rectal swab | 17.72 | no variants | no variants |
| | RM8 | BAL | 80.01 | no variants | no variants |
| | | LLLL | 16.3 | no variants | no variants |
| | | Nose swab | 0 | ND | ND |
| | | RLLL | 4.28 | ND | ND |
| | | Rectal swab | 1.39 | ND | no variants |

| | | | | |
|---|---|---|---|---|
| RM9 | BAL | 258.39 | no variants | no variants |
| | LLLL | 1.34 | ND | no variants |
| | Nose swab | 0.87 | ND | ND |
| | RLLL | 26.65 | no variants | no variants |
| RM10 | BAL | 250.71 | no variants | no variants |
| | LLLL | 7.12 | no variants | no variants |
| | Nose swab | 0.2 | ND | ND |
| | RLLL | 1210.31 | no variants | no variants |
| | Rectal swab | 5.17 | ND | no variants |
| RM11 | BAL | 880.97 | no variants | no variants |
| | LLLL | 56.38 | no variants | no variants |
| | Nose swab | 2.63 | ND | ND |
| | RLLL | 597.65 | no variants | no variants |
| | Rectal swab | 0.1 | ND | ND |
| RM12 | BAL | 415.3 | no variants | no variants |
| | LLLL | 0.43 | ND | ND |
| | Nose swab | 20.44 | no variants | no variants |
| | RLLL | 88.08 | no variants | no variants |
| | Rectal swab | 11.56 | ND | no variants |

[a] BAL: bronchoalveolar lavages collected at 3 dpi.

[b] LLLL: left lower lung lobe collected on 7 dpi

[c] collected on 5 dpi

[d] RLLL: right lower lung lobe collected on 7 dpi

[e] collected on 2 dpi

[f] No sequence coverage or coverage was too limited to call

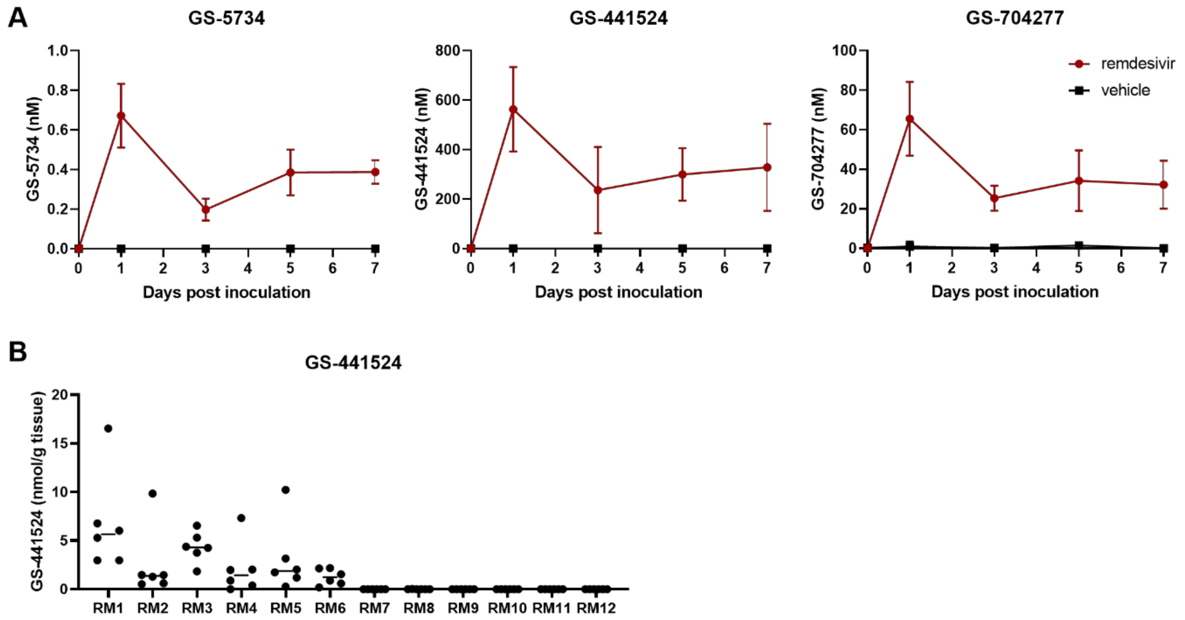


**Figure S1. Concentration of remdesivir prodrug and metabolites measured in serum and lung homogenates of rhesus macaques infected with SARS-CoV-2.** Two groups of six rhesus macaques were inoculated with SARS-CoV-2 strain nCoV-WA1-2020. Twelve hours post inoculation, one group was administered 10mg/kg intravenous remdesivir and the other group was treated with an equal volume of vehicle solution (2ml/kg). Treatment was continued 12hrs after the first treatment, and every 24 hrs thereafter with a dose of 5 mg/kg remdesivir or equal volume of vehicle solution (1ml/kg). Panel A shows the serum concentration of remdesivir prodrug GS-5734, the dephosphorylated nucleoside product GS-441524 and the intermediate alanine metabolite GS-704277 over time as measured in by LCMS. Mean and standard deviation are shown. Panel B shows the concentration of GS-441524 homogenized lung tissue collected from all six lung lobes on 7 dpi, 24 hrs after the last remdesivir treatment was administered. Each dot represents the concentration of GS-441524 in one lung lobe.
